# A novel metabarcoded DNA sequencing tool for the detection of *Plasmodium* species in malaria positive patients

**DOI:** 10.1101/801175

**Authors:** Abdul Wahab, Ayaz Shaukat, Qasim Ali, Mubashir Hussain, Taj Ali Khan, M. Azmat Ullah Khan, Imran Rashid, Mushtaq A. Saleem, Mike Evans, Neil D. Sargison, Umer Chaudhry

## Abstract

Various PCR based methods have been described for the diagnosis of malaria, but most depend on the use of *Plasmodium* species-specific probes and primers; hence only the tested species are identified and there is limited available data on the true circulating species diversity. Sensitive diagnostic tools and platforms for their use are needed to detect *Plasmodium* species in both clinical cases and asymptomatic infections that contribute to disease transmission. We have been recently developed for the first time a novel high throughput ‘haemoprotobiome’ metabarcoded DNA sequencing method and applied it for the quantification of haemoprotozoan parasites (*Theleria* and *Babesia*) of livestock. Here, we describe a novel, high throughput method using an Illumina MiSeq platform to demonstrate the proportions of *Plasmodium* species in metabarcoded DNA samples derived from human malaria patients. *Plasmodium falciparum* and *Plasmodium vivax* positive control gDNA was used to prepare mock DNA pools of parasites to evaluate the detection threshold of the assay for each of the two species and to assess the accuracy of proportional quantification. We then applied the assay to malaria-positive human samples to show the species composition of *Plasmodium* communities in the Punjab province of Pakistan and in the Afghanistan-Pakistan tribal areas. The diagnostic performance of the deep amplicon sequencing method was compared to an immunochromatographic assay that is widely used in the region. Metabarcoded DNA sequencing showed better diagnostic performance, greatly increasing the estimated prevalence of *Plasmodium* infection. The next-generation sequencing method using metabarcoded DNA has potential applications in the diagnosis, surveillance, treatment, and control of *Plasmodium* infections, as well as to study the parasite biology.

## 1. Introduction

Malaria is the most important vector-borne disease, causing high morbidity and mortality (Moody, 2002). More than 3.4 billion people are infected, resulting in an estimated 1.2 billion malaria cases every year (Poostchi et al., 2018). The disease is caused by intracellular protozoan parasites of the genus *Plasmodium* transmitted through *Anopheles* mosquitoes (Dash et al., 2007; Sinka et al., 2012). *Plasmodium* has an indirect life cycle including one stage in *Anopheles* mosquitoes and three different stages in humans, all with different rates of replication (Li et al., 1997). Gametogenesis occurs in human blood and fertilisation of male and female macro and microgametes occurs in the midgut of the mosquito after feeding. Asexual stages occur in the gut of the mosquito as sporogony, and after biting the humans, sporozoites undergo exoerythrocytic schizogeny in the hepatic cells and then erythrocytic schizogony in the blood cells (Li et al., 1994b). Five species of *Plasmodium* parasite infect humans, namely *Plasmodium falciparum*, *Plasmodium vivax*, *Plasmodium ovale*, *Plasmodium malariae* and *Plasmodium knowlesi*. *P. falciparum* is the most associated with lethal disease, but in recent years, there has been an increase in disease severity attributable to *P. vivax* (Saravu et al., 2014; Scholzen et al., 2014; William et al., 2011).

The 18S rDNA of *Plasmodium* is unique due to its genomic arrangement dispersed among different chromosomes. The copy number is limited to 4 to 8 per genome and the sequences are not identical (Li et al., 1994b). Their expression is regulated during different development stages of the life cycle (Li et al., 2014); for example, at least three types of genotypic variants have been identified between stages of *P. falciparum* and *P. vivax* laboratory isolates (Li et al., 1994b; McCutchan et al., 1988; Qari et al., 1994; Rogers et al., 1995). However, the presence of these variants has not been reported in the field studies.

The 18S rDNA of *Plasmodium* forms a mosaic of conserved and variable regions; whereby the conserved regions contribute to form a secondary structure of rRNA that appears to be associated with the universal function of the ribosomes. The variable regions are scattered among the conserved regions and contribute to major differences in gene composition and size (Li et al., 1994a). The function of variable regions is not fully understood, but determination of sequence variations can discriminate between *Plasmodium* species (Agudelo et al., 2013; Haanshuus et al., 2013; Lee et al., 2015; Lefterova et al., 2015), and overcome limitations of traditional microscopic and immunochromatographic methods for the diagnosis of this group of parasites at species level.

Significant progress has been made in the global fight against malaria through high throughput rapid diagnosis. Sensitive diagnostic tools are needed to detect clinically and subclinically infected patients (Echeverry et al., 2016). Molecular methods including qPCR, species-specific PCR, nested PCR, and multiplex PCR have been described (Canier et al., 2013; Cunha et al., 2009; Das et al., 1995; Echeverry et al., 2016; Haanshuus et al., 2013; Steenkeste et al., 2009), but these are low throughput, hence relatively expensive (Chaudhry et al., 2019). These methods depend on the use of species-specific probes and primers, meaning that only the tested species are identified, hence are limited in their ability to describe true circulating species diversity (Moody, 2002). In contrast, high throughput metabarcoded DNA sequencing using the Illumina MiSeq platform is relatively low-cost and potentially less error-prone. We have applied this ‘haemprotobiome’ method, to the study of tick-borne haemoprotozoan parasites of ruminants (Chaudhry et al., 2019). The method has the potential to open new areas of research in the study of *Plasmodium*, to accurately provide relative quantification of co-infecting species and to evaluate drug treatment responses (Shaukat et al., 2019). The method uses primers binding to the conserved sites and analyse of up to 600 bp sequence reads. The use of adapter and barcoded primers allows a large number of samples to be pooled and sequenced in a single MiSeq flow cell, making the assay suitable for high-throughput analysis (Shaukat et al., 2019).

Here, we report for the first time the development of a deep sequencing method using the Illumina MiSeq platform to quantify *P. falciparum* and *P. vivax* present in malaria-positive human blood samples. The results are compared with a standard immunochromatographic assay to validate the method’s accuracy for species identification.

## 2. Materials and methods

### 2.1. Parasite material

Positive control samples of *P. falciparum* were kindly provided by D Jason Mooney at the Roslin Institute, University of Edinburgh, UK, previously obtained from National Institute for Biological Standards and Control (NIBSC code 04/176). *P. vivax* control samples were kindly provided by Dr Imran Rashid at the University of Veterinary and Animal Sciences, Lahore, Pakistan, previously obtained from the Biodefense and Emerging Infections Research Resources Repository (BEI code MRA-41). Four replicates each of mock pools comprising of *P. falciparum* only (Mix 1), *P. vivax* only (Mix 2), and *P. falciparum* and *P. vivax* (Mix 3) were created. These were used to test the detection threshold of the metabarcoded sequencing method and to show the proportions of each of the *Plasmodium* species present. Four negative control of human blood samples were provided by Sana Amir and Saqib Shahzad, Chughtai Diagnostic Laboratory, Lahore Pakistan.

Malaria suspected patients referred to Basic Health Units in the tribal areas of the Afghanistan-Pakistan border and Chughtai Diagnostic Laboratory in the Punjab province of Pakistan were invited to participate in this study. Prior discussions were held with key administrative and community leaders to raise awareness of the study. Samples were taken by trained para-medical workers under the supervision of local collaborators and the Basic Health Unit or Chughtai Diagnostic Laboratory staff. The institutional review boards of the University of Central Punjab (UCP-30818), and the Kohat University of Science and Technology, Pakistan (KUST/EC/1379) approved the study. Patients of all age groups were included in this study with symptoms consistent with malaria, including vomiting, fever, headache, chills, sweats, nausea and fatigue.

Blood samples were collected by venipuncture during peak malaria transmission seasons between August to November 2017 and 2018. A total of 5 ml of intravenous blood was drawn into EDTA tubes and stored at −20 ^0^C for gDNA isolation. Each sample was also routinely analysed by microscopic examination under oil immersion (x1000) of 4% Giemsa-stained blood smears for the diagnosis of malaria. The *Plasmodium* goes through different stages of their development cycle (48 hr), which gives the parasites a different visual appearance that can be observed under the microscope. These stages show the ring (Fig. 1A), tropnozoite, schizont, and gametocyte appearance. Malaria case identification was based on the appearance of those stages on the microscopic examination (Fig. 1A). Overall, 365 malaria suspected positive patients were identified.

**Fig. 1.**
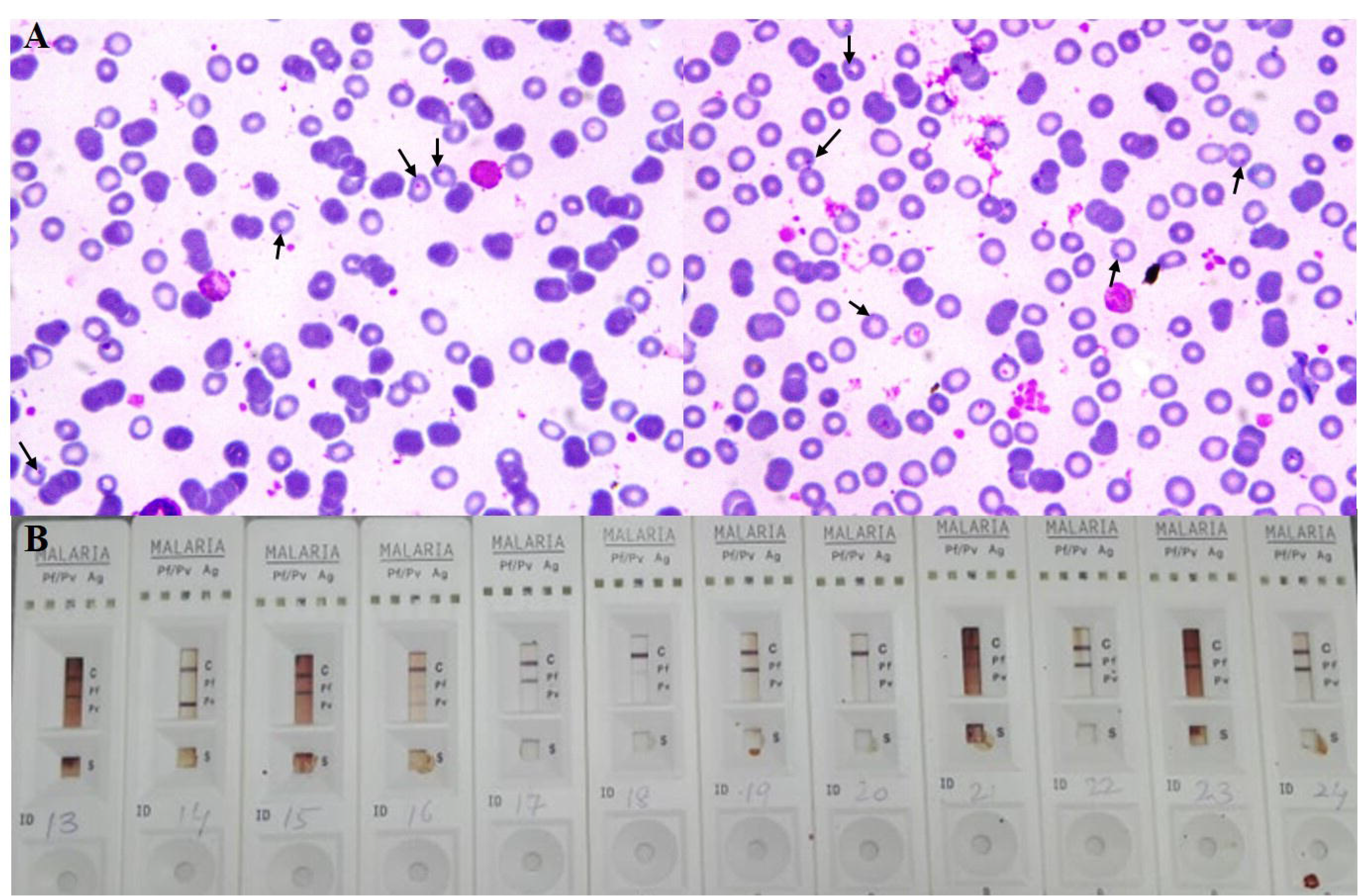
(A) Giemsa-stained blood smears were examined by 1000 x microscopy, showing rings in the *Plasmodium* positive samples. (B) Immunochromatographic assay for the detection of *P. falciparum* specific histidine-rich protein 2 (Pf-HRP2) and *P. vivax* specific lactate dehydrogenase (Pv-LDH). Adequate volumes of the blood samples were dispensed into the sample well ‘S’ of the test cassette. If Pf-HRP2 was bind to the HRP2 gold conjugates forming a burgundy colored pf band, indicating *P. falciparum* positive test. If Pv-LDH was bind to the LDH gold conjugates forming a burgundy colored pv band, indicating *P. vivax* positive test. The absence of any band suggests a negative result and C is positive control.

### 2.2. Immunochromatographic assay

The 365 malaria-positive on microscopic examination blood samples, each having an unknown level of parasitemia, were analysed using a commercial immunochromatographic rapid diagnostic test (RDT) kit. The Malaria Pf/Pv Ag Rapid Test (Healgen®; Zhejiang Orient Gene Biotech Co, Ltd) RDT kit was designed to detect *P. falciparum*-specific histidine-rich protein 2 (Pf-HRP2) and *P.vivax*-specific lactate dehydrogenase (Pv-LDH). The kit was transported and maintained at the room temperature, opened just before use to avoid humidity damage, and used in accordance with the manufacturer’s recommendations. During the assay, an adequate volume of the blood sample was dispensed into the sample well ‘S’ of the test cassette and the lysis buffer is added to the buffer well ‘B’. The buffer contains a detergent that lyses the red blood cells and releases antigens, which migrate by capillary action across the strip held in the cassette. If Pf-HRP2 binds to the HRP2 gold conjugates and the immunocomplex is then captured on the membrane by the pre-coated anti-Pf-HRP2 antibodies, forming a burgundy colored pf band, indicating *P. falciparum* positive test (Fig. 1B). If Pv-LDH binds to the LDH gold conjugates and the immunocomplex is then captured on the membrane by the pre-coated anti-Pv-LDH antibodies, forming a burgundy colored pv band, indicating *P. vivax* positive test (Fig. 1B). The absence of any band suggests a negative result. The test also contained an internal control ‘C’ band, exhibiting a burgundy colored band of the immunocomplex of goat anti-mouse IgG/mouse IgG (anti-Pv-LDH and anti-Pf-HRP2) gold conjugates, regardless of the color development on ‘C’ band.

### 2.3. Genomic DNA isolation, primer design and adapter/barcoded PCR amplification of rDNA 18S locus

50 μl of blood from each of the 365 samples was used as a template, to extract gDNA according to the protocols described in the TIANamp blood DNA kit (Tiangen Biotech (Beijing) Co., Ltd). Overall, 589 bp and 568 bp fragments of *P. falciparum* and *P. vivax* 18S rDNA, respectively, were amplified using newly developed adapter primer sets (Supplementary Table S1). The overall scheme of the sample preparation is described in Figure 2A. Adapters were added to these primers to allow the successive annealing of subsequent metabarcode primers and N is the number of random nucleotides included between the locus-specific primers and adapter to increase the variety of generated amplicons as previously described by Chaudhry et al. (2019). Four forward (Plasmo1_For, Plasmo1_For-1N, Plasmo1_For-2N, Plasmo1_For-3N) and four reverse (Plasmo2_Rev, Plasmo1_Rev-1N, Plasmo1_Rev-2N, Plasmo1_Rev-3N) primers were mixed in equal proportion (Supplementary Table S1) and used for first-round PCR under the following conditions: 5X KAPA buffer, 10mM dNTPs, 10 uM forward and reverse adapter primer, 0.5 U KAPA Polymerase (KAPA Biosystems, USA), and 1 ul of worm lysate. The thermocycling conditions of the PCR were 95°C for 2 min, followed by 35 cycles of 98°C for 20 sec, 60°C for 15 sec, 72°C for 15 sec and a final extension 72°C for 5 min. PCR products were purified with AMPure XP Magnetic Beads (1X) (Beckman Coulter, Inc.).

**Fig. 2.**
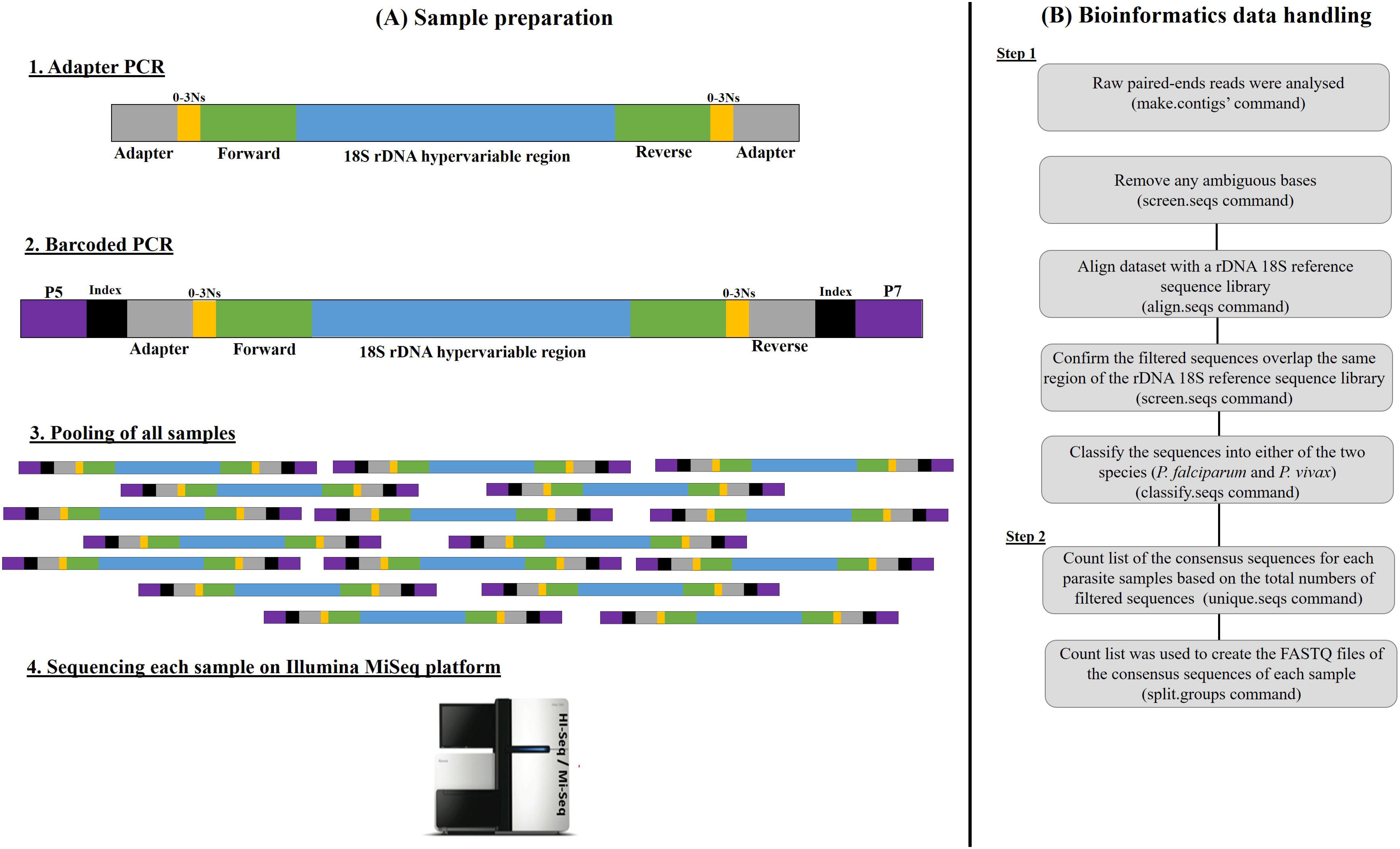
Schematic representation of the sample preparation (A) and the bioinformatics data handling (B) of the metabarcoded sequencing library. (A) In the first-round PCR amplification, overhanging forward and reverse primers were used to amplify the rDNA 18S. The adapter base pairs provide the target sites for the primers used for sequencing and the random nucleotides (0-3Ns) were inserted between the primers and the adapter to offset the reading frame, therefore amplicons prevent the oversaturation of the MiSeq sequencing channels. The second-round PCR amplification was then performed using overhanging barcoded primers bound to the adapter tags to add indices, as well as the P7 and P5 regions required to bind to the MiSeq flow cell. (B) Text files containing rDNA 18S sequence data (FASTQ files) were generated from the Illumina MiSeq binary raw data outputs, and data analyses were performed using a bespoke modified pipeline in Mothur v1.39.5 software (Schloss et al., 2009) and Illumina MiSeq standard procedures (Kozich et al., 2013) as described in materials and methods section 2.4.

After the purification, a second-round PCR was performed by using sixteen forward and twenty-four reverse barcoded primers. The barcoded forward (N501 to N516) and reverse (N701 to N724) primers (10 uM each) were previously described by Chaudhry et al. (2019). The primers were used in a manner that repetition of same forward and reverse sequences did not occur in the different samples. The second-round PCR conditions were: 5X KAPA buffer, 10 mM dNTPs 0.5 U KAPA Polymerase (KAPA Biosystems, USA), and 2 ul of first-round PCR product as DNA template. The thermocycling conditions of the second round PCR were 98°C for 45 sec, followed by 7 cycles of 98°C for 20 sec, 63°C for 20 sec, and 72°C for 2 minutes. PCR products were purified with AMPure XP Magnetic Beads (1X) according to the protocols described by Beckman Coulter, Inc.

### 2.4. Sequencing of metabarcoded 18S rDNA, data handling, and bioinformatic analysis

The pooled library was measured with KAPA qPCR library quantification kit (KAPA Biosystems, USA). The prepared library was then run on Illumina MiSeq Sequencer using a 600-cycle pair-end reagent kit (MiSeq Reagent Kits v2, MS-103-2003) at a concentration of 15 nM with an addition of 25% Phix Control v3 (Illumina, FC-11-2003).

The overall scheme of the data handling and bioinformatics analysis is described in Figure 2B. MiSeq data were handled with our own bioinformatics pipeline (Chaudhry et al., 2019). Briefly, MiSeq separates all sequence reads during post-run processing using the barcoded indices and to generate FASTQ files. The raw paired read-ends were run into the ‘make.contigs’ command to combine the two sets of reads for each sample. The command extracts sequence and quality score data from the FASTQ files, creating the complement of the reverse and forward reads, and then joining the reads into contigs. After removing the too long, or ambiguous sequence reads, the data were then aligned with the *P. falciparum* and *P. vivax* reference sequence library (for more details Supplementary Data S1 and Result section 3.1) using the ‘align.seqs’ command to summarise the 589 bp and 568 bp fragments encompassing parts of the 18S DNA spanning the hyper-variable region of the *P. falciparum* and *P. vivax* ribosomal cistrons. At this stage, the 18S rDNA analysis was completed by classifying the sequences into either of the two species by using the ‘classify.seqs’ command and creating a taxonomy file by using the ‘summary.tax’ command. Overall, 762674 million reads of 18S rDNA were generated from the data set.

For the phylogenetic analysis of *P. falciparum* and *P. vivax* 18S rDNA, all the classified sequences were run on the ‘screen.seqs’ command and the count list of the consensus sequences of each sample was created using the ‘unique.seqs’ command followed by the use of the ‘pre.cluster’ command to look for sequences differences and to merge them in groups based on their abundance. Any chimeras were identified and removed by using the ‘chimera.vsearch’ command. The count list was further used to create the FASTQ files of the consensus sequences of each sample using the ‘split.groups’ command (for more details Supplementary Data S2).

### 2.5. *Split and maximum-likelihood trees of P. falciparum and* P. vivax *18S rDNA sequences*

*P. falciparum* and *P. vivax* 18S rDNA sequence reads were analysed separately in Geneious v9.0.1 software (Kearse et al., 2012) using the MUSCLE alignment tool. The aligned sequences were then imported into the FaBox 1.5 online tool to collapse those with 100% base pair similarity after corrections into single genotypes. The split tree of *P. falciparum* and *P. vivax* 18S rDNA was created in the SplitTrees4 software by using the UPGMA method in the Jukes-Cantor model of substitution (Huson and Bryant, 2005). The maximum-likelihood tree for *P. falciparum* and *P. vivax* 18S rDNA was constructed by the HKY model of substitution in the MEGA 7 software and to select the appropriate model of nucleotide substitutions (Tamura et al., 2013).

### 2.6. Statistical analysis

The data were analysed using CompareTests: Correct for Verification Bias in Diagnostic Accuracy and Agreement Statistics software (R package version 1.2.). The frequency of *P. falciparum* and *P. vivax* in the samples was calculated by dividing the number of sequence reads for each sample by the total number of reads. The effect of the mock pools of *P. falciparum* and *P. vivax* positive controls were analysed by running a Kruskal-Wallis rank-sum test for each admixture. The performance of the immunochromatographic assay was compared against metabarcoded sequencing, using a Fisher’s Exact test to calculate the predictive value and Category-Specific Classification Probability (CSCP) with 95% confidence interval. Kappa (k) values were calculated to express the agreement beyond chance; where values greater than 0.80 were considered to represent perfect agreement; values of 0.61 - 0.80 to represent good agreement; and values of 0.21 - 0.60 to represent moderate agreement.

## 3. Results

### 3.1. *P. falciparum and P.* vivax *18S rDNA reference sequence libraries*

The sequence reads generated by metabarcoded 18S rDNA sequencing of the *P. falciparum* and *P. vivax* positive controls were compared to the malaria-positive samples and to the published NCBI GenBank 18S rDNA sequences to account for any genetic diversity. A total of 34 *P. falciparum* and 74 *P. vivax* reference sequences were identified (Fig. 3, Supplementary Data S1). The maximum-likelihood tree shows distinct clustering of the *P. falciparum* and *P. vivax* 18S rDNA regions (Fig. 3), hence these closely related *Plasmodium* species can be reliably differentiated by virtue of 18S rDNA sequence variations.

**Fig. 3.**
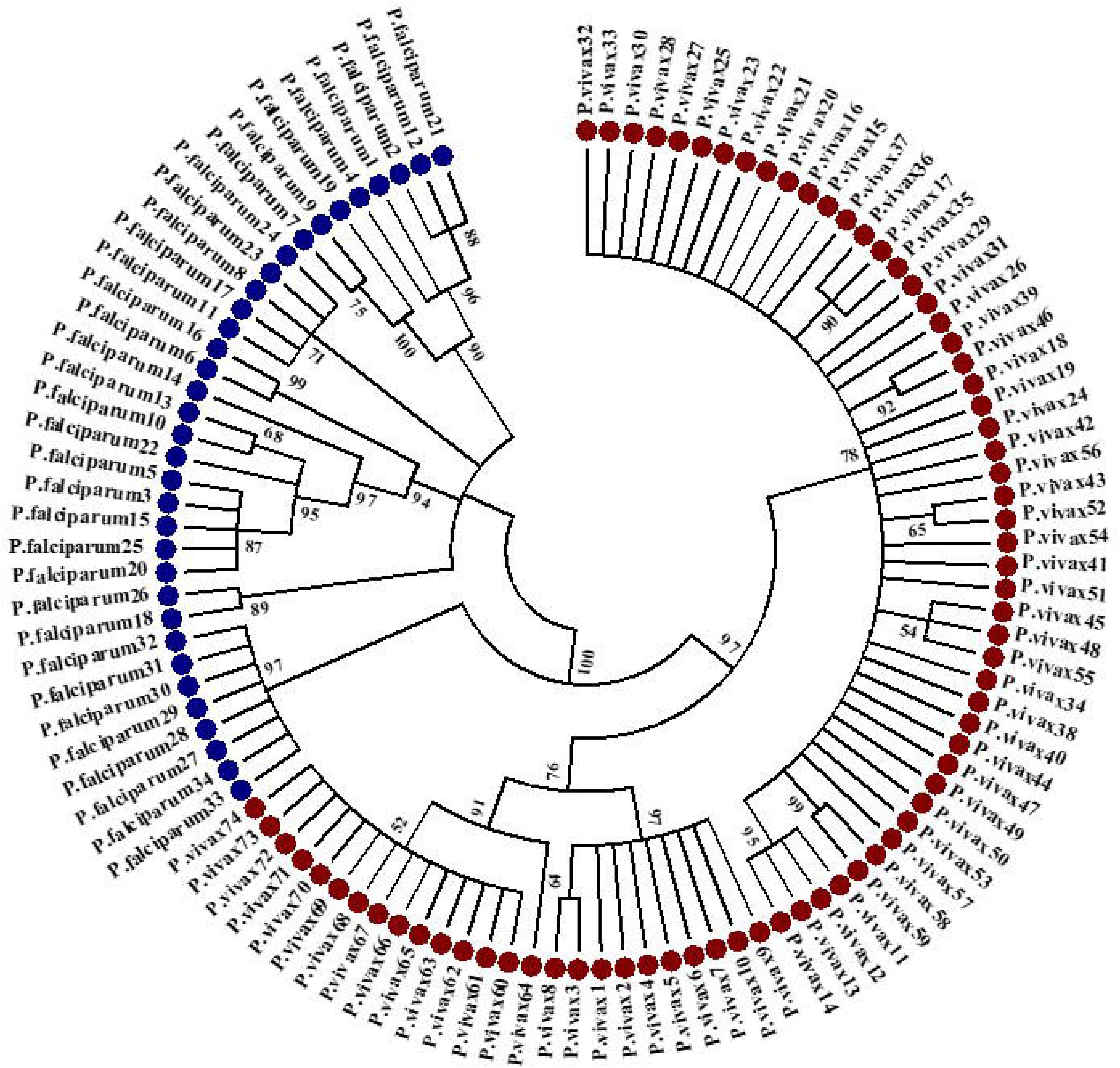
The maximum-likelihood tree was obtained from the *P. falciparum* and *P. vivax* rDNA 18S region. The sequences were first calculated the number of reference sequences generated from both species (Supplementary Data S1). A total of 34 reference sequences of the rDNA 18S locus were identified in *P. falciparum* and 74 reference sequences were identified in *P. vivax*. The reference sequences were aligned on the MUSCLE tool of the Geneious v9.0.1 software. The neighbor-joining algorithm (HKY parameter model) was computed with 1000 bootstrap replicates using MEGA 7 software. Both species were identified with different color shades (*P. falciparum* in blue and *P. vivax* in brown).

### 3.2. Validation of the metabarcoded sequencing assay using mock pools of P. falciparum and P. vivax

Four replicates each of *P. falciparum* only (Mix 1), *P. vivax* only (Mix 2), and *P. falciparum* and *P. vivax* (Mix 3) were created from gDNA to demonstrate the detection accuracy of the metabarcoded DNA sequencing method and to show the proportions of each of the species being present (Fig. 4; Supplementary Table S2). The mixing of different mock pools demonstrates the accurate detection ability of the metabarcoded sequencing method and to show the proportions of each of the species being present. The Mix 1 pool yielded only *P. falciparum* sequence reads and the Mix 2 pool yielded only *P. vivax* sequence reads (Fig. 4). The Mix 3 pool yielded both *P. falciparum* and *P. vivax* sequence reads, with no statistically significant variations between replicates (Kruskal-Wallis rank-sum test; Mix1: χ^2^(1) 0, p=1; Mix2: χ^2^(1) 0, p=1; Mix3: χ^2^(3) 0.02153, p=0.5231).

**Fig. 4.**
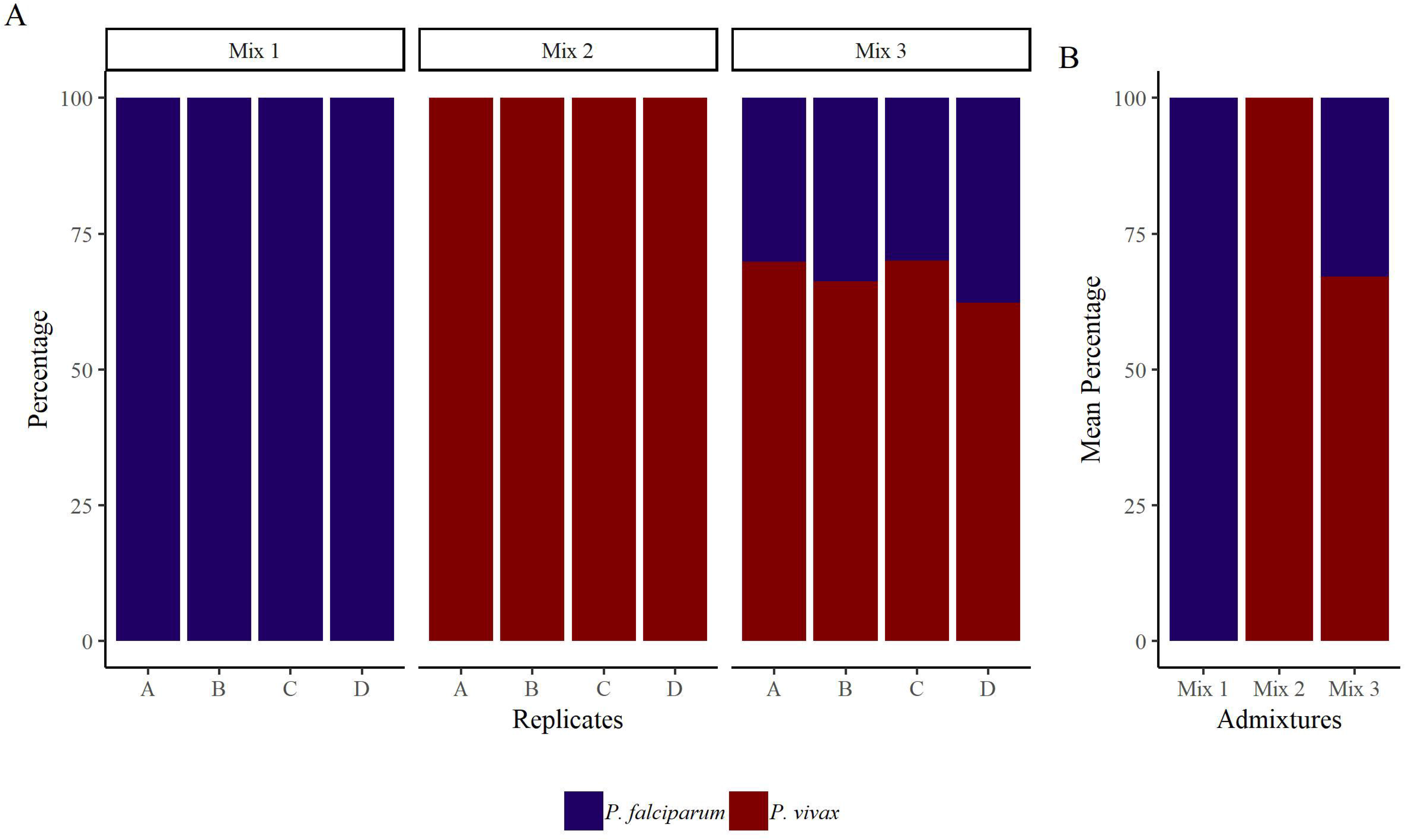
Validation of the metabarcoding sequencing using three separate mock pools [Mix 1 (*P. falciparum*), Mix 2 (*P. vivax*), Mix 3 (*P. falciparum* and *P. vivax*)] of unknown numbers of parasites from each species. Panel 2A shows that metabarcoded sequencing was used on four replicates of each mock pool to amplify both species as denoted on the X-axis. The Y-axis shows the percentage proportions of each species. Panel 2B shows how the replicates were grouped and averaged based on the amplification. The species are identified with different colours (*P. falciparum* in blue and *P. vivax* in brown).

### 3.3. Assessment of the immunochromatographic assay and metabarcoded sequencing for the identification of P. falciparum and P. vivax

The immunochromatographic assay and metabarcoded DNA sequencing methods were applied to malaria-positive blood samples to detect *P. falciparum* and *P. vivax* in the field (Fig. 5, Supplementary Table S3). The results of both assays demonstrate that the prevalence of *P. vivax* infection was higher than that of *P. falciparum* infection. In the case of the metabarcoded DNA sequencing assay, those samples yielding more than 1000 reads (implying sufficient gDNA for accurate amplification) were included in the analysis (Supplementary Table S3). *Plasmodium vivax* was present in 199 (69.8%) patients, *P. falciparum* in 84 (29.5%) and mixed infection in 2 (0.7%) patients (Fig. 5). The immunochromatographic assay showed that *Plasmodium vivax* was present in 187 (65.6%) patients, *P. falciparum* in 78 (27.4%), mixed infection in 2 (0.7%) patients (Fig. 5) and 18 (6.32%) malaria-positive cases were negative in RDT, but positive in the metabarcoded DNA sequencing assay (Fig. 5).

**Fig. 5.**
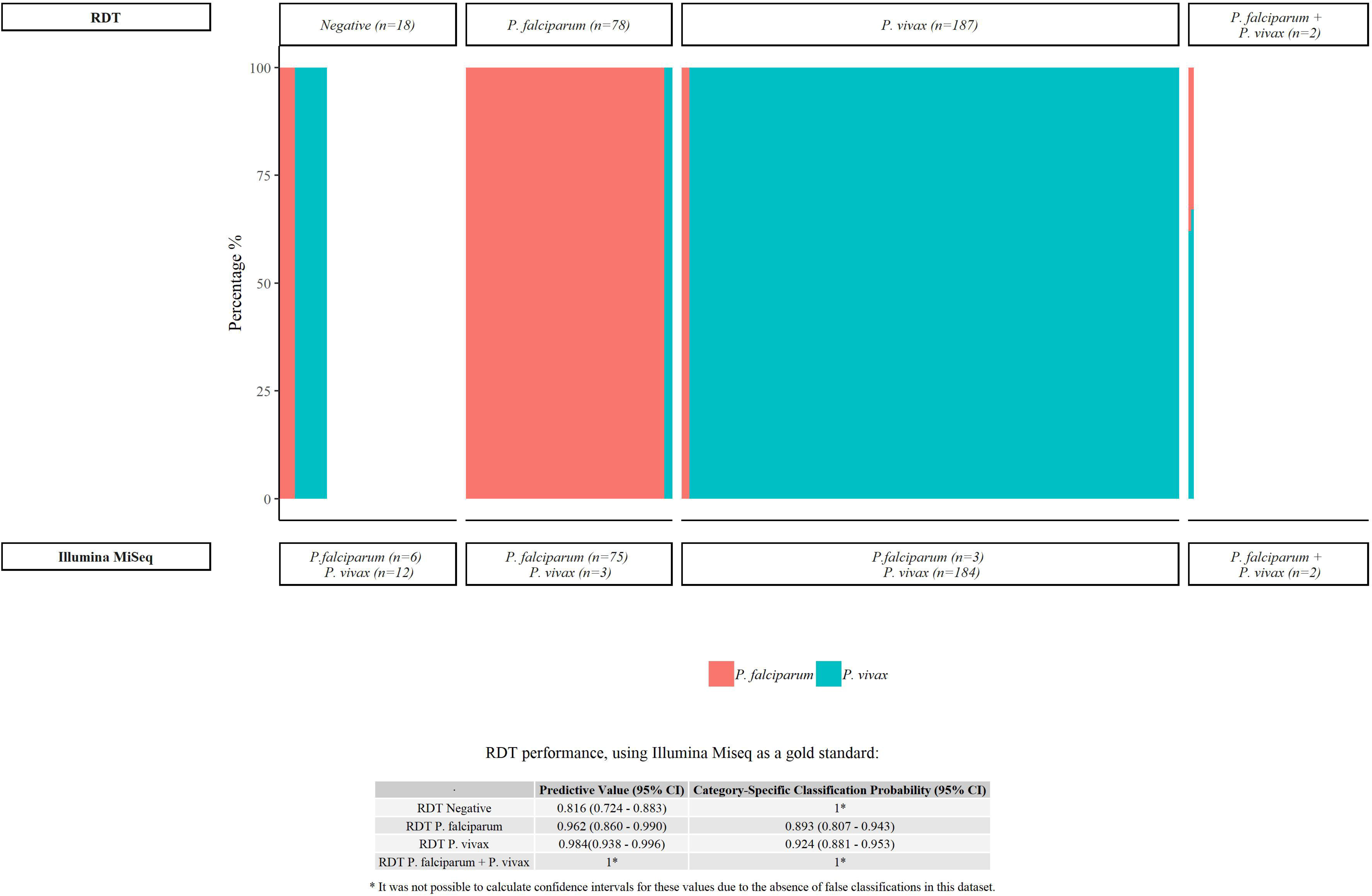
Immunochromatographic assay (RDT) and the metabarcoding sequencing (Illumina MiSeq) was performed for the detection of *P. falciparum* and *P. vivax*. A total of 365 malaria-positive samples were collected in EDTA tube from basic health units in the tribal area of the Pakistan-Afghanistan border and Chughtai Diagnostic Laboratory in the Punjab province of Pakistan. The immunochromatographic assay and the metabarcoded 18S rDNA sequencing methods were applied to each sample; the X-axis shows the proportion of each species being estimated and the Y-axis shows the percentage proportions of each species. The species are identified with different colours (*P. falciparum* in pink and *P. vivax* in light blue).

The degree of agreement between the immunochromatographic and metabarcoded DNA sequencing assays was high with κ = 0.893 (95% CI: 0.839-0.930). The Category-Specific Classification Probability (CSCP) was also high in all four categories, ranging from 0.892 (95% CI: 0.807 - 0.943) to 1. The Predictive Values (PV) for ‘positive RDT’ results were also very high, ranging from 0.962 (0.860 - 0.990) to 1. However, the PVs for ‘negative RDT’ results were 0.816 (0.724 - 0.883); being significantly lower than the Predictive Value for both ‘RDT *P. falciparum*’ (p=0.004) and ‘RDT *P. vivax*’ (p<0.001) results. In samples with similar disease prevalence, there is, therefore, an increased likelihood that a negative RDT result may be incorrect.

### 3.4. Phylogenetic analysis of the P. falciparum and P. vivax rDNA 18S sequences

Overall, 112 different genotypes of the *P. falciparum* 18S rDNA locus were identified among 84 field samples and 12 from the NCBI GenBank sequences (Supplementary Data S2 and section 3.1. and 3.3). The split tree shows at least two distinct clades (Fig. 6A), and it sets apart that 9 genotypes in the Clade I, belongs to a ‘type S’ 18S rDNA region (McCutchan et al., 1988). The remaining 115 genotypes were in clade II, belonging to a ‘type A’ 18S rDNA region (McCutchan et al., 1988). 80 genotypes predominated in group 1 and 35 genotypes in group 2 (Fig. 6A). In contrast, 30 different genotypes of the *P. vivax* 18S rDNA locus was identified among 199 field samples and 18 from the NCBI GenBank sequences (Supplementary Data S2 and section 3.1. and 3.3). The split tree shows at least two distinct clades (Fig. 6B), and it sets apart that 3 genotypes in the Clade I, belong to the ‘type S’ 18S rDNA as described by Li et al. (1994). The remaining 45 genotypes were in clade II, belonging to ‘type A’ 18S rDNA (Li et al., 1994b). 7 genotypes predominated in groups 1 and 2, and 31 genotypes in group 3 (Fig. 6B).

**Fig. 6.**
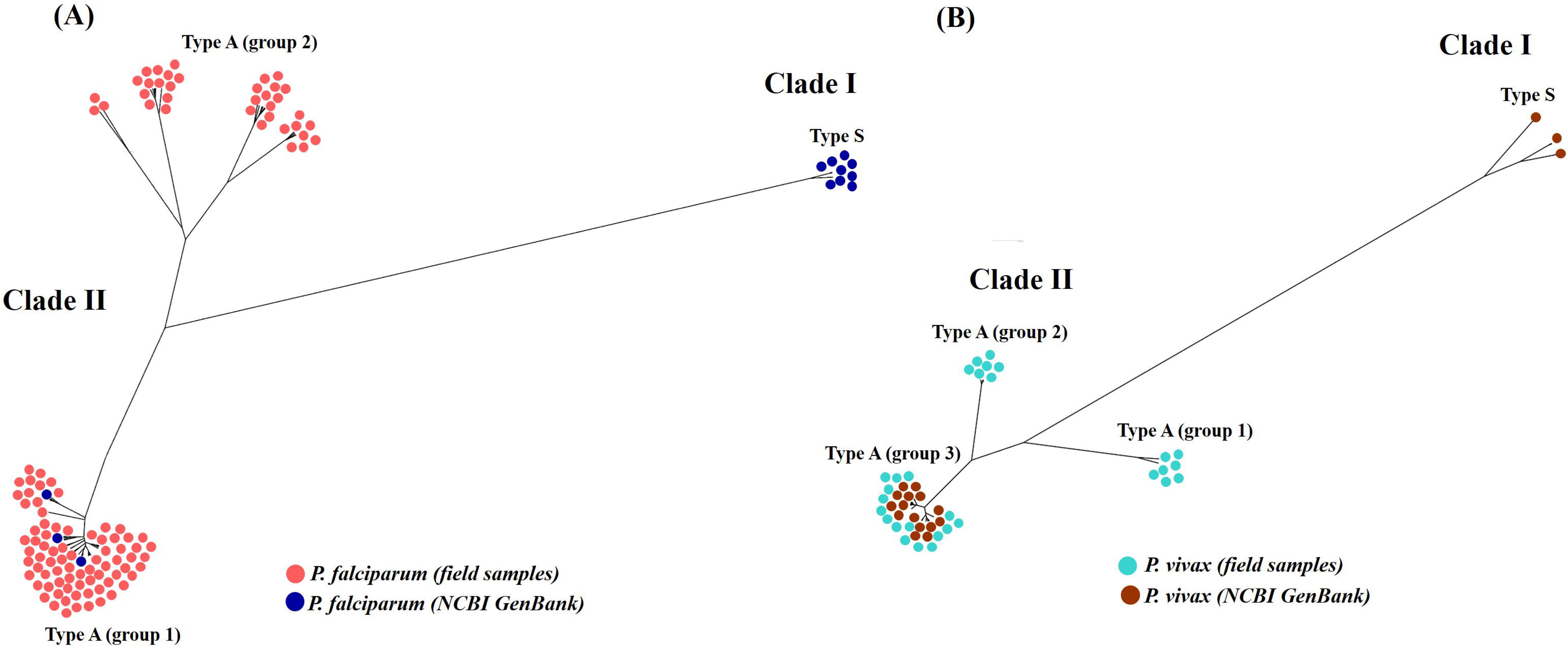
Split tree was made for the *P. falciparum* and *P. vivax* 18S rDNA sequence data. 134 genotypes were identified in *P. falciparum* and 48 genotypes were identified in *P. vivax* (Supplementary Data S2). The genotypes were aligned on the MUSCLE tool of the Geneious v9.0.1 and the tree was constructed with the UPGMA method in the Jukes-Cantor model of substitution in the SplitsTrees4 software. The appropriate model of nucleotide substitutions was selected by using the jModeltest 13.1.0 program. The pie chart circles in the tree represent the different 18S genotypes containing different colours as follows: (A) *P. falciparum* from the field samples are coloured pink (type A group 1 and 2) and NCBI database sequences are coloured blue (type A group 1 and type S). (B) *P. vivax* from the field samples are coloured light blue (type A group 1, 2 and 3) and NCBI database sequences are coloured brown (type A group 3 and type S).

## 4. Discussion

Microscopic examination of blood smears remains the mainstay for the diagnosis of *Plasmodium* in the field. The procedure allows the detection of levels of at least 200 parasites/ul, which is sufficient for the diagnosis of most symptomatic cases, but can result in misdiagnoses at low levels of parasitemia (Rakotonirina et al., 2008; Wongsrichanalai et al., 2007). The method is labor-intensive, time-consuming, and interpretation of results requires highly skilled microscopists (Canier et al., 2013; Echeverry et al., 2016).

The immunochromatographic assay is also utilised in case investigation and malaria surveillance programs in the field. The method depends on the incorporation of conjugated monoclonal antibodies providing the indicator of infection (Moody, 2002; Wongsrichanalai et al., 2007). The targeted antigens are abundant in the sexual and asexual stages of the parasites, while histidine-rich protein 2 (Pf-HRP2) based RDT is specific for *P. falciparum* and lactate dehydrogenase (Pf-LDH & Pv-LDH) specific for detecting both *P. falciparum* and *P. vivax*. Pan based RDT targets specific antigens including lactate dehydrogenase (P-LDH) and aldolase proteins found in all *Plasmodium* species (Akinyi Okoth et al., 2015; Murillo Solano et al., 2015). Several factors potentially affect the accuracy and false-negative results of RDT including the interpretation of the test strip colour change, the density of malaria infection in the host, improper storage/handling of the kit and poor test performance (Echeverry et al., 2016). Besides this, other major factors such cross-reactivity of HRP2 with histidine-rich protein 3 (a structural homolog with significant sequence similarity) and deletions in the HRP2 locus in *P. falciparum* isolates may account for false-negative results (Akinyi Okoth et al., 2015; Rachid Viana et al., 2017).

Conventional PCR based molecular methods are useful in the detection of *Plasmodium* species for which the reagents and conditions have been developed, but have limitations in terms of lacking scalability (Canier et al., 2013; Cunha et al., 2009; Das et al., 1995; Echeverry et al., 2016; Haanshuus et al., 2013; Steenkeste et al., 2009). The diagnostic challenges of the disease identification have not been resolved yet (Wongsrichanalai et al., 2007), therefore the metabarcod DNA sequencing potentially provides a more accurate and reliable automated high-throughput method to detect *Plasmodium* species in blood samples. The use of a single PCR utilising primers conserved between *Plasmodium* species provides a powerful tool to measure the relative sequence representation of each species in the blood samples. In the present study, we have evaluated a metabarcoded DNA sequencing method to identify the presence of *P. falciparum* and *P. vivax* using mock parasite pools, applying the method to malaria-positive blood samples, and the detection *P. falciparum* and *P. vivax* rDNA18S genotypic variants in the field samples.

We tested the ability of the metabarcoded DNA sequencing assay to accurately determine the relative species proportions in combinations of *P. falciparum* and *P. vivax*. To do this, we generated mock pools containing different estimated proportions of both species and demonstrated no significant variations between replicates. A previous study using pools of laboratory-maintained *Theileria* and *Babesia* haemoprotozoan parasites showed that the relative sequence representation was unaffected by either the number of PCR cycles employed or the parasite species composition of the sample. This study also found no sequence representation bias in PCR products used for sequencing, arising from the number of first-round PCR cycles (Chaudhry et al., 2019).

After validating the metabarcoded DNA sequencing assay using mock pools of *Plasmodium* positive DNA, we applied the method to field samples collected from suspected malaria-positive patients in the tribal area of the Afghanistan-Pakistan border and in the Punjab province of Pakistan, where malaria caused by *P. falciparum* and *P. vivax* has been reported (Kakar et al., 2010; Khattak et al., 2013). Our findings support previous reports of the high prevalence of malaria in the tribal areas of the Afghanistan-Pakistan border, and increasing prevalence over the last few decades in the Punjab province (Kakar et al., 2010; Khattak et al., 2013). Our results support the reports suggesting that while the majority of the cases of malaria are caused by *P. vivax,* the prevalence of *P. falciparum* has increased during recent years (Khattak et al., 2013). The increased prevalence of *P. falciparum* may be an attribute to antimicrobial resistance; previous studies have shown that the pyrimethamine and chloroquine resistance mutations in *P. falciparum* are present in different cities of Pakistan (Ghanchi et al., 2011). Another explanation for the increased prevalence of *P. falciparum* may be provided by the influx of people and the movement of refugees from areas of Afganistan where the parasite species is common (Howard et al., 2011).

Three structurally distinct types of rDNA 18S genotypic variants have been reported in *P. falciparum* and *P. vivax* laboratory isolates (Li et al., 1994b; McCutchan et al., 1988; Qari et al., 1994; Rogers et al., 1995). The existence of genotypic variants in the field studies has not been described. In the present study, we have identified the two independent gene duplication events that occurred in *P. falciparum*, leading to the A and S type rDNA18S lineages; the type A lineage being the ancestor of at least two groups (Fig. 6A). In *P. vivax*, we identified two independent gene duplication events, also leading to the A and S type rDNA18S lineages; the type A lineage being the ancestor of at least three groups (Fig. 6B). In the previous reports, type A was transcribed in erythrocytic schizogony and gametocyte stages consistent with those stages that could have been represented in the present study. In these reports, type S was transcribed in the exoerythrocytic schizogeny stage, while type O was associated with oocyst development only in infected mosquitoes (Li et al., 1997). These observed differences in the 18S loci of *P. falciparum* and *P. vivax* field samples confirm the presence of genotypic variants. Better understanding is needed of the function of these structurally distinct ribosomes that are active with enhanced transcription during different stages of parasitic development in *Plasmodium,* with reference to the development of disease control strategies.

In conclusion, we describe for the first time the use of metabarcoded DNA sequencing using an Illumina MiSeq platform to quantify *P. falciparum* and *P. vivax,* and demonstrate its accuracy on malaria-positive samples. Our results provide a proof of concept for the use of the method in disease surveillance, similar to its application in the study of haemoprotozoan parasites of livestock (Chaudhry et al., 2019) and dogs (Huggins et al., 2019). This work was undertaken to explore the possibilities for the application of this high throughput method to determine the dynamics of co-infections, disease biology and epidemiology in *Plasmodium* parasites, and has applications in monitoring the changes in parasite populations after the emergence and spread of antimicrobial drug resistance (Shaukat et al., 2019).

## Acknowledgment

The study was financially supported by the Roslin Institute uses facilities funded by the Biotechnology and Biological Sciences Research Council (BBSRC). Work at the University of Veterinary and Animal Science Pakistan and Kohat University of Science and Technology Pakistan uses facilities funded by the Higher Education Commission of Pakistan.

## Conflict of interest

None

## Notes

#### Summary of Updates

Various PCR based methods have been described for the diagnosis of malaria but most depend on the use of Plasmodium species specific probes and primers; hence only the tested species are identified and there is limited available data on the true circulating species diversity. Sensitive diagnostic tools and platforms for their use are needed to detect Plasmodium species in both clinical cases and asymptomatic infections that contribute to disease transmission. We have been recently developed for the first time a novel high throughput haemoprotobiome metabarcoded DNA sequencing method and applied it for the quantification of haemoprotozoan parasites (Theleria and Babesia) of livestock. Here we describe a novel, high throughput method using an Illumina MiSeq platform to demonstrate the proportions of Plasmodium species in metabarcoded DNA samples derived from human malaria patients. Plasmodium falciparum and Plasmodium vivax positive control gDNA was used to prepare mock DNA pools of parasites to evaluate the detection threshold of the assay for each of the two species and to assess the accuracy of proportional quantification. We then applied the assay to malaria-positive human samples to show the species composition of Plasmodium communities in the Punjab province of Pakistan and in the Afghanistan-Pakistan tribal areas. The diagnostic performance of the deep amplicon sequencing method was compared to an immunochromatographic assay that is widely used in the region. Metabarcoded DNA sequencing showed better diagnostic performance, greatly increasing the estimated prevalence of Plasmodium infection. The next-generation sequencing method using metabarcoded DNA has potential applications in the diagnosis, surveillance, treatment, and control of Plasmodium infections, as well as to study the parasite biology.

