## Supplementary Table S1 for "A novel metabarcoded DNA sequencing tool for the detection of *Plasmodium* species in malaria positive patients"

**Supplementary Table S1.** Primer sequences for Illumina MiSeq Library preparation. Plasmo 1-For / Plasmo 2-Rev primer sequence are underlined, N’s are bolded.

| **Sequences (5'-3')** | **Primer Name** |
| --- | --- |
| TCGTCGGCAGCGTCAGATGTGTATAAGAGACAG*GAGGTAGTGACAAGAAATAACAA*T*A* | Plasmo 1-For |
| TCGTCGGCAGCGTCAGATGTGTATAAGAGACAGN*GAGGTAGTGACAAGAAATAACAA*T*A* | Plasmo 1-For1N |
| TCGTCGGCAGCGTCAGATGTGTATAAGAGACAGNN*GAGGTAGTGACAAGAAATAACAA*T*A* | Plasmo 1-For2N |
| TCGTCGGCAGCGTCAGATGTGTATAAGAGACAGNNN*GAGGTAGTGACAAGAAATAACAA*T*A* | Plasmo 1-For3N |
| GTCTCGTGGGCTCGGAGATGTGTATAAGAGACAG*TATCTGATCGTCTTCACTCCCTTA*A*C* | Plasmo 2-Rev |
| GTCTCGTGGGCTCGGAGATGTGTATAAGAGACAGN*TATCTGATCGTCTTCACTCCCTTA*A*C* | Plasmo 2-Rev1N |
| GTCTCGTGGGCTCGGAGATGTGTATAAGAGACAGNN*TATCTGATCGTCTTCACTCCCTTA*A*C* | Plasmo 2-Rev2N |
| GTCTCGTGGGCTCGGAGATGTGTATAAGAGACAGNNN*TATCTGATCGTCTTCACTCCCTTA*A*C* | Plasmo 2-Rev3N |
