## Supplementary Table S2 for "A novel metabarcoded DNA sequencing tool for the detection of *Plasmodium* species in malaria positive patients"

| **Sample** | **Mix** | **Replicate** | ***P. falciparum*** | ***P. vivax*** | **Total** |
| --- | --- | --- | --- | --- | --- |
| M1A | 1 | A | 8615 | 0 | 8615 |
| M1B | 1 | B | 9711 | 0 | 9711 |
| M1C | 1 | C | 7922 | 0 | 7922 |
| M1D | 1 | D | 8881 | 0 | 8881 |
| M2A | 2 | A | 0 | 12689 | 35129 |
| M2B | 2 | B | 0 | 13889 | 13889 |
| M2C | 2 | C | 0 | 11403 | 11403 |
| M2D | 2 | D | 0 | 14220 | 14220 |
| M3A | 3 | A | 5246 | 12129 | 17375 |
| M3B | 3 | B | 6814 | 13335 | 20149 |
| M3C | 3 | C | 5741 | 13451 | 19192 |
| M3D | 3 | D | 6636 | 10964 | 17600 |
|  |  |  | 24437 | 49879 | 74316 |

**Supplementary Table S2.** The mock pools were made from *P. falciparum* and *P. vivax* with random number of parasites from each species, to test the threshold of the metabarcoding sequencing for each of the species.
