## Supplementary Table S3 for "A novel metabarcoded DNA sequencing tool for the detection of *Plasmodium* species in malaria positive patients"

**Supplementary Table S3.** *Plasmodium* species have been identified by microscopy, rapid diagnostic test and Illumina MiSeq from the blood samples collected from different cities of the Tribal Areas of Pakistan-Afganistan border and the Punjab province of Pakistan.

| **Sample ID** | **City** | **Province/border area** | **Microscopy** | **Rapid Diagnostic Test** | **Illumina MiSeq** | |
| --- | --- | --- | --- | --- | --- | --- |
|  |  |  |  |  | ***P. falciparum*** | ***P. vivax*** |
| 1 | Lahore | Punjab | *Plasmodium* | *P.vivax* |  | 2145 |
| 2 | Lahore | Punjab | *Plasmodium* | *P.vivax* |  | 3102 |
| 3 | Lahore | Punjab | *Plasmodium* | *P.vivax* |  | 1895 |
| 4 | Lahore | Punjab | *Plasmodium* | *P.vivax* |  | 3439 |
| 5 | Lahore | Punjab | *Plasmodium* | *P.vivax* |  | 4194 |
| 6 | Sheikhupura | Punjab | *Plasmodium* | *P.vivax* |  | 2271 |
| 7 | Sheikhupura | Punjab | *Plasmodium* | *P.vivax* |  | 2105 |
| 8 | Sheikhupura | Punjab | *Plasmodium* | *P.vivax* |  | 1549 |
| 9 | Sheikhupura | Punjab | *Plasmodium* | *P.vivax* |  | 1213 |
| 10 | Gujranwala | Punjab | *Plasmodium* | *P.vivax* |  | 2167 |
| 11 | Gujrat | Punjab | *Plasmodium* | *P.vivax* |  | 3482 |
| 12 | Lahore | Punjab | *Plasmodium* | *P.vivax* |  | 1132 |
| 13 | Lahore | Punjab | *Plasmodium* | *P.vivax* |  | 3245 |
| 14 | Lahore | Punjab | *Plasmodium* | *P.vivax* |  | 2283 |
| 15 | Lahore | Punjab | *Plasmodium* | *P.vivax* |  | 2675 |
| 16 | Lahore | Punjab | *Plasmodium* | *P.vivax* |  | 2132 |
| 17 | Lahore | Punjab | *Plasmodium* | *P.vivax* |  | 3015 |
| 18 | Narowal | Punjab | *Plasmodium* | *P.vivax* |  | 2213 |
| 19 | Lahore | Punjab | *Plasmodium* | *P.vivax* |  | 3743 |
| 20 | Lahore | Punjab | *Plasmodium* | *P.vivax* |  | 3590 |
| 21 | Lahore | Punjab | *Plasmodium* | *P.vivax* |  | 2923 |
| 22 | Lahore | Punjab | *Plasmodium* | *P.vivax* |  | 2113 |
| 23 | Lahore | Punjab | *Plasmodium* | *P.vivax* |  | 1147 |
| 24 | Lahore | Punjab | *Plasmodium* | *P.vivax* |  | 1748 |
| 25 | Multan | Punjab | *Plasmodium* | *P.vivax* |  | 1516 |
| 26 | Lahore | Punjab | *Plasmodium* | *P.vivax* |  | 2818 |
| 27 | Lahore | Punjab | *Plasmodium* | *P.vivax* |  | 1269 |
| 28 | Lahore | Punjab | *Plasmodium* | *P.vivax* |  | 2417 |
| 29 | Gujranwala | Punjab | *Plasmodium* | *P.vivax* |  | 2864 |
| 30 | Faisalabad | Punjab | *Plasmodium* | *P.vivax* |  | 1932 |
| 31 | Lahore | Punjab | *Plasmodium* | *P.vivax* |  | 2826 |
| 32 | Lahore | Punjab | *Plasmodium* | *P.vivax* |  | 4755 |
| 33 | Multan | Punjab | *Plasmodium* | *P.vivax* |  | 3043 |
| 34 | Lahore | Punjab | *Plasmodium* | *P.vivax* |  | 4822 |
| 35 | Gujrat | Punjab | *Plasmodium* | *P.vivax* |  | 2617 |
| 36 | Lahore | Punjab | *Plasmodium* | *P.vivax* |  | 4220 |
| 37 | D.G. Khan | Punjab | *Plasmodium* | *P.vivax* |  | 2033 |
| 38 | Lahore | Punjab | *Plasmodium* | *P.vivax* |  | 4063 |
| 39 | Lahore | Punjab | *Plasmodium* | *P.vivax* |  | 3156 |
| 40 | Lahore | Punjab | *Plasmodium* | *P.vivax* |  | 2139 |
| 41 | Lahore | Punjab | *Plasmodium* | *P.vivax* |  | 3521 |
| 42 | Lahore | Punjab | *Plasmodium* | *P.vivax* |  | 3444 |
| 43 | Lahore | Punjab | *Plasmodium* | *P.vivax* |  | 4145 |
| 44 | Lahore | Punjab | *Plasmodium* | *P.vivax* |  | 4917 |
| 45 | Lahore | Punjab | *Plasmodium* | *P.vivax* |  | 3299 |
| 46 | Lahore | Punjab | *Plasmodium* | *P.vivax* |  | 2100 |
| 47 | Lahore | Punjab | *Plasmodium* | *P.vivax* |  | 2794 |
| 48 | Lahore | Punjab | *Plasmodium* | *P.vivax* |  | 4104 |
| 49 | Lahore | Punjab | *Plasmodium* | *P.vivax* |  | 2598 |
| 50 | D.G.Khan | Punjab | *Plasmodium* | *P.vivax* |  | 3897 |
| 51 | Lahore | Punjab | *Plasmodium* | *P.vivax* |  | 4112 |
| 52 | Lahore | Punjab | *Plasmodium* | *P.vivax* |  | 3678 |
| 53 | Lahore | Punjab | *Plasmodium* | *P.vivax* |  | 2073 |
| 54 | Lahore | Punjab | *Plasmodium* | *P.vivax* |  | 3767 |
| 55 | Lahore | Punjab | *Plasmodium* | *P.vivax* |  | 4761 |
| 56 | Lahore | Punjab | *Plasmodium* | *P.vivax* |  | 2635 |
| 57 | Multan | Punjab | *Plasmodium* | *P.vivax* |  | 3483 |
| 58 | Bhakkar | Punjab | *Plasmodium* | *P.vivax* |  | 3161 |
| 59 | Lahore | Punjab | *Plasmodium* | *P.vivax* |  | 2137 |
| 60 | Lahore | Punjab | *Plasmodium* | *P.vivax* |  | 2110 |
| 61 | Lahore | Punjab | *Plasmodium* | *P.vivax* |  | 1129 |
| 62 | D.G.Khan | Punjab | *Plasmodium* | *P.vivax* |  | 4113 |
| 63 | Lahore | Punjab | *Plasmodium* | *P.vivax* |  | 3104 |
| 64 | Lahore | Punjab | *Plasmodium* | *P.vivax* |  | 4737 |
| 65 | Gujranwala | Punjab | *Plasmodium* | *P.vivax* |  | 3149 |
| 66 | Lahore | Punjab | *Plasmodium* | *P.vivax* |  | 2539 |
| 67 | Lahore | Punjab | *Plasmodium* | *P.vivax* |  | 3452 |
| 68 | Multan | Punjab | *Plasmodium* | *P.vivax* |  | 2168 |
| 69 | Lahore | Punjab | *Plasmodium* | *P.vivax* |  | 3175 |
| 70 | Sheikhupura | Punjab | *Plasmodium* | *P.vivax* |  | 2764 |
| 71 | Lahore | Punjab | *Plasmodium* | *P.vivax* |  | 4131 |
| 72 | Multan | Punjab | *Plasmodium* | *P.vivax* |  | 3155 |
| 73 | Lahore | Punjab | *Plasmodium* | *P.vivax* |  | 4266 |
| 74 | Lahore | Punjab | *Plasmodium* | *P.vivax* |  | 3207 |
| 75 | Lahore | Punjab | *Plasmodium* | *P.vivax* |  | 4123 |
| 76 | Lahore | Punjab | *Plasmodium* | *P.vivax* |  | 2208 |
| 77 | Lahore | Punjab | *Plasmodium* | *P.vivax* |  | 3149 |
| 78 | Lahore | Punjab | *Plasmodium* | *P.vivax* |  | 4222 |
| 79 | Lahore | Punjab | *Plasmodium* | *P.vivax* |  | 5112 |
| 80 | Lahore | Punjab | *Plasmodium* | *P.vivax* |  | 3435 |
| 81 | Lahore | Punjab | *Plasmodium* | *P.vivax* |  | 4226 |
| 82 | Lahore | Punjab | *Plasmodium* | *P.vivax* |  | 2178 |
| 83 | Multan | Punjab | *Plasmodium* | *P.vivax* |  | 3169 |
| 84 | Lahore | Punjab | *Plasmodium* | *P.vivax* |  | 4110 |
| 85 | Lahore | Punjab | *Plasmodium* | *P.vivax* |  | 1164 |
| 86 | Lahore | Punjab | *Plasmodium* | *P.vivax* |  | 3129 |
| 87 | D.G.Khan | Punjab | *Plasmodium* | *P.vivax* |  | 2693 |
| 88 | Lahore | Punjab | *Plasmodium* | *P.vivax* |  | 2132 |
| 89 | Rahem Yar Khan | Punjab | *Plasmodium* | *P.vivax* |  | 1146 |
| 90 | Rahem Yar Khan | Punjab | *Plasmodium* | *P.vivax* |  | 2105 |
| 91 | Lahore | Punjab | *Plasmodium* | *P.vivax* |  | 4330 |
| 92 | Lahore | Punjab | *Plasmodium* | *P.vivax* |  | 3121 |
| 93 | Lahore | Punjab | *Plasmodium* | *P.vivax* |  | 4177 |
| 94 | Lahore | Punjab | *Plasmodium* | *P.vivax* |  | 4352 |
| 95 | Lahore | Punjab | *Plasmodium* | *P.vivax* |  | 1216 |
| 96 | D.G. Khan | Punjab | *Plasmodium* | *P.vivax* |  | 2115 |
| 97 | Lahore | Punjab | *Plasmodium* | *P.vivax* |  | 2100 |
| 98 | Lahore | Punjab | *Plasmodium* | *P.vivax* |  | 1986 |
| 99 | Lahore | Punjab | *Plasmodium* | *P.vivax* |  | 1654 |
| 100 | Lahore | Punjab | *Plasmodium* | *P.vivax* |  | 2156 |
| 101 | Lahore | Punjab | *Plasmodium* | *P.vivax* |  | 1521 |
| 102 | Lahore | Punjab | *Plasmodium* | *P.vivax* |  | 1973 |
| 103 | Lahore | Punjab | *Plasmodium* | *P.vivax* |  | 2100 |
| 104 | Lahore | Punjab | *Plasmodium* | *P.vivax* |  | 2346 |
| 105 | Lahore | Punjab | *Plasmodium* | *P.vivax* |  | 3881 |
| 106 | Lahore | Punjab | *Plasmodium* | *P.vivax* |  | 4160 |
| 107 | Lahore | Punjab | *Plasmodium* | *P.vivax* |  | 2826 |
| 108 | Lahore | Punjab | *Plasmodium* | *P.vivax* |  | 3534 |
| 109 | Lahore | Punjab | *Plasmodium* | *P.vivax* |  | 3230 |
| 110 | Lahore | Punjab | *Plasmodium* | *P.vivax* |  | 4148 |
| 111 | Lahore | Punjab | *Plasmodium* | *P.vivax* |  | 2625 |
| 112 | Lahore | Punjab | *Plasmodium* | *P.vivax* |  | 1872 |
| 113 | Lahore | Punjab | *Plasmodium* | *P.vivax* |  | 2919 |
| 114 | Lahore | Punjab | *Plasmodium* | *P.vivax* |  | 2106 |
| 115 | Lahore | Punjab | *Plasmodium* | *P.vivax* |  | 2815 |
| 116 | Lahore | Punjab | *Plasmodium* | *P.vivax* |  | 3224 |
| 117 | Lahore | Punjab | *Plasmodium* | *P.vivax* |  | 3213 |
| 118 | Lahore | Punjab | *Plasmodium* | *P.vivax* |  | 3228 |
| 119 | Lahore | Punjab | *Plasmodium* | *P.vivax* |  | 2185 |
| 120 | Lahore | Punjab | *Plasmodium* | *P.vivax* |  | 4109 |
| 121 | Lahore | Punjab | *Plasmodium* | *P.vivax* |  | 2341 |
| 122 | Lahore | Punjab | *Plasmodium* | *P.vivax* |  | 3227 |
| 123 | Lahore | Punjab | *Plasmodium* | *P.vivax* |  | 3110 |
| 124 | Lahore | Punjab | *Plasmodium* | *P.vivax* |  | 3289 |
| 125 | Lahore | Punjab | *Plasmodium* | *P.vivax* |  | 1918 |
| 126 | Lahore | Punjab | *Plasmodium* | *P.vivax* |  | 1119 |
| 127 | Lahore | Punjab | *Plasmodium* | *P.vivax* |  | 1290 |
| 128 | Lahore | Punjab | *Plasmodium* | *P.vivax* |  | 2611 |
| 129 | Lahore | Punjab | *Plasmodium* | *P.vivax* |  | 3759 |
| 130 | Lahore | Punjab | *Plasmodium* | *P.vivax* |  | 3353 |
| 131 | Gujranwala | Punjab | *Plasmodium* | *P.vivax* |  | 3183 |
| 132 | Multan | Punjab | *Plasmodium* | *P.vivax* |  | 2339 |
| 133 | Faisalabad | Punjab | *Plasmodium* | *P.vivax* |  | 1424 |
| 134 | Sheikhopora | Punjab | *Plasmodium* | *P.vivax* |  | 1852 |
| 135 | Sarghodah | Punjab | *Plasmodium* | *P.vivax* |  | 1271 |
| 136 | Sheikhopora | Punjab | *Plasmodium* | *P.vivax* |  | 3138 |
| 137 | Lahore | Punjab | *Plasmodium* | *P.vivax* |  | 4277 |
| 138 | karachi | Punjab | *Plasmodium* | *P.vivax* |  | 1197 |
| 139 | karachi | Punjab | *Plasmodium* | *P.vivax* |  | 1248 |
| 140 | Lahore | Punjab | *Plasmodium* | *P.vivax* |  | 1258 |
| 141 | Lahore | Punjab | *Plasmodium* | *P.vivax* |  | 2626 |
| 142 | Lahore | Punjab | *Plasmodium* | *P.vivax* |  | 3220 |
| 143 | DG Khan | Punjab | *Plasmodium* | *P.vivax* |  | 3234 |
| 144 | Sheikhopora | Punjab | *Plasmodium* | *P.vivax* |  | 2212 |
| 145 | Faisalabad | Punjab | *Plasmodium* | *P.vivax* |  | 1155 |
| 146 | Gujrat | Punjab | *Plasmodium* | *P.vivax* |  | 2263 |
| 147 | Gujranwala | Punjab | *Plasmodium* | *P.vivax* |  | 4354 |
| 148 | Sheikhupura | Punjab | *Plasmodium* | *P.vivax* |  | 1259 |
| 149 | DG Khan | Punjab | *Plasmodium* | *P.vivax* |  | 2118 |
| 150 | Lahore | Punjab | *Plasmodium* | *P.vivax* |  | 2158 |
| 151 | Lahore | Punjab | *Plasmodium* | *P.vivax* |  | 3292 |
| 152 | Lahore | Punjab | *Plasmodium* | *P.vivax* |  | 2340 |
| 153 | Lahore | Punjab | *Plasmodium* | *P.vivax* |  | 3270 |
| 154 | Lahore | Punjab | *Plasmodium* | *P.vivax* |  | 3577 |
| 155 | Lahore | Punjab | *Plasmodium* | *P.vivax* |  | 2131 |
| 156 | Lahore | Punjab | *Plasmodium* | *P.vivax* |  | 3261 |
| 157 | Lahore | Punjab | *Plasmodium* | *P.vivax* |  | 2237 |
| 158 | Lahore | Punjab | *Plasmodium* | *P.vivax* |  | 2384 |
| 159 | Kasur | Punjab | *Plasmodium* | *P.vivax* |  | 3122 |
| 160 | Lahore | Punjab | *Plasmodium* | *P.vivax* |  | 3219 |
| 161 | Shaikhupora | Punjab | *Plasmodium* | *P.vivax* |  | 3132 |
| 162 | Kasur | Punjab | *Plasmodium* | *P.vivax* |  | 2330 |
| 163 | Lahore | Punjab | *Plasmodium* | *P.vivax* |  | 2323 |
| 164 | Lahore | Punjab | *Plasmodium* | *P.vivax* |  | 3311 |
| 165 | Lahore | Punjab | *Plasmodium* | *P.vivax* |  | 2119 |
| 166 | Lahore | Punjab | *Plasmodium* | *P.vivax* |  | 2483 |
| 167 | Lahore | Punjab | *Plasmodium* | *P.vivax* |  | 3489 |
| 168 | Shaikhupora | Punjab | *Plasmodium* | *P.vivax* |  | 3230 |
| 169 | Kasur | Punjab |  | *P.vivax* |  | 2352 |
| 170 | Lahore | Punjab | *Plasmodium* | *P.vivax* |  | 3441 |
| 171 | Lahore | Punjab | *Plasmodium* | *P.vivax* |  | 3175 |
| 172 | Shaikhupora | Punjab | *Plasmodium* | *P.vivax* |  | 3364 |
| 173 | Lahore | Punjab | *Plasmodium* | *P.vivax* |  | 3459 |
| 174 | Lahore | Punjab | *Plasmodium* | *P.vivax* |  | 3214 |
| 175 | Lahore | Punjab | *Plasmodium* | *P.vivax* |  | 2198 |
| 176 | Shaikhupora | Punjab | *Plasmodium* | *P.vivax* |  | 3668 |
| 177 | Lahore | Punjab | *Plasmodium* | *P.vivax* |  | 1261 |
| 178 | Multan | Punjab | *Plasmodium* | *P.vivax* |  | 1301 |
| 179 | Lahore | Punjab | *Plasmodium* | *P.vivax* |  | 3103 |
| 180 | Gujranwala | Punjab | *Plasmodium* | *P.vivax* |  | 3182 |
| 181 | Lahore | Punjab | *Plasmodium* | *P.vivax* |  | 2100 |
| 182 | Lahore | Punjab | *Plasmodium* | *P.vivax* |  | 2316 |
| 183 | Multan | Punjab | *Plasmodium* | *P.vivax* |  | 2548 |
| 184 | Shanaweri Zagiri | Tribal Area (Orakzai Agency) | *Plasmodium* | *P.vivax* |  | 2212 |
| 185 | Lahore | Punjab | *Plasmodium* | *P.falciparum* | 2723 |  |
| 186 | D.G.Khan | Punjab | *Plasmodium* | *P.falciparum* | 2294 |  |
| 187 | Lahore | Punjab | *Plasmodium* | *P.falciparum* | 3929 |  |
| 188 | Multan | Punjab | *Plasmodium* | *P.falciparum* | 1408 |  |
| 189 | Lahore | Punjab | *Plasmodium* | *P.falciparum* | 1015 |  |
| 190 | D.G.Khan | Punjab | *Plasmodium* | *P.falciparum* | 1455 |  |
| 191 | D.G.Khan | Punjab | *Plasmodium* | *P.falciparum* | 4692 |  |
| 192 | Shanaweri Zagiri | Tribal Area (Orakzai Agency) | *Plasmodium* | *P.falciparum* | 2762 |  |
| 193 | Shanaweri Zagiri | Tribal Area (Orakzai Agency) | *Plasmodium* | *P.falciparum* | 2567 |  |
| 194 | Shanaweri Zagiri | Tribal Area (Orakzai Agency) | *Plasmodium* | *P.falciparum* | 1870 |  |
| 195 | Shanaweri Zagiri | Tribal Area (Orakzai Agency) | *Plasmodium* | *P.falciparum* | 1267 |  |
| 196 | Shanaweri Zagiri | Tribal Area (Orakzai Agency) | *Plasmodium* | *P.falciparum* | 1383 |  |
| 197 | Shanaweri Zagiri | Tribal Area (Orakzai Agency) | *Plasmodium* | *P.falciparum* | 1468 |  |
| 198 | Shanaweri Zagiri | Tribal Area (Orakzai Agency) | *Plasmodium* | *P.falciparum* | 1386 |  |
| 199 | Shanaweri Zagiri | Tribal Area (Orakzai Agency) | *Plasmodium* | *P.falciparum* | 1762 |  |
| 200 | Shanaweri Zagiri | Tribal Area (Orakzai Agency) | *Plasmodium* | *P.falciparum* | 1319 |  |
| 201 | Ali Masjid | Tribal Area (Khyber Agency) | *Plasmodium* | *P.falciparum* | 2312 |  |
| 202 | Ali Masjid | Tribal Area (Khyber Agency) | *Plasmodium* | *P.falciparum* | 2191 |  |
| 203 | Ali Masjid | Tribal Area (Khyber Agency) | *Plasmodium* | *P.falciparum* | 2912 |  |
| 204 | Ali Masjid | Tribal Area (Khyber Agency) | *Plasmodium* | *P.falciparum* | 1635 |  |
| 205 | Ali Masjid | Tribal Area (Khyber Agency) | *Plasmodium* | *P.falciparum* | 2029 |  |
| 206 | Ali Masjid | Tribal Area (Khyber Agency) | *Plasmodium* | *P.falciparum* | 2535 |  |
| 207 | Ali Masjid | Tribal Area (Khyber Agency) | *Plasmodium* | *P.falciparum* | 1821 |  |
| 208 | Ali Masjid | Tribal Area (Khyber Agency) | *Plasmodium* | *P.falciparum* | 1616 |  |
| 209 | Ali Masjid | Tribal Area (Khyber Agency) | *Plasmodium* | *P.falciparum* | 2614 |  |
| 210 | Ali Masjid | Tribal Area (Khyber Agency) | *Plasmodium* | *P.falciparum* | 1197 |  |
| 211 | Ali Masjid | Tribal Area (Khyber Agency) | *Plasmodium* | *P.falciparum* | 2378 |  |
| 212 | Ali Masjid | Tribal Area (Khyber Agency) | *Plasmodium* | *P.falciparum* | 2768 |  |
| 213 | Ali Masjid | Tribal Area (Khyber Agency) | *Plasmodium* | *P.falciparum* | 2225 |  |
| 214 | Ali Masjid | Tribal Area (Khyber Agency) | *Plasmodium* | *P.falciparum* | 2359 |  |
| 215 | Ali Masjid | Tribal Area (Khyber Agency) | *Plasmodium* | *P.falciparum* | 3427 |  |
| 216 | Ali Masjid | Tribal Area (Khyber Agency) | *Plasmodium* | *P.falciparum* | 2949 |  |
| 217 | Ali Masjid | Tribal Area (Khyber Agency) | *Plasmodium* | *P.falciparum* | 3263 |  |
| 218 | Ali Masjid | Tribal Area (Khyber Agency) | *Plasmodium* | *P.falciparum* | 2306 |  |
| 219 | Ali Masjid | Tribal Area (Khyber Agency) | *Plasmodium* | *P.falciparum* | 1319 |  |
| 220 | Ali Masjid | Tribal Area (Khyber Agency) | *Plasmodium* | *P.falciparum* | 1121 |  |
| 221 | Ali Masjid | Tribal Area (Khyber Agency) | *Plasmodium* | *P.falciparum* | 3880 |  |
| 222 | Ali Masjid | Tribal Area (Khyber Agency) | *Plasmodium* | *P.falciparum* | 2990 |  |
| 223 | Ali Masjid | Tribal Area (Khyber Agency) | *Plasmodium* | *P.falciparum* | 3173 |  |
| 224 | Ali Masjid | Tribal Area (Khyber Agency) | *Plasmodium* | *P.falciparum* | 1275 |  |
| 225 | Ali Masjid | Tribal Area (Khyber Agency) | *Plasmodium* | *P.falciparum* | 2456 |  |
| 226 | Ali Masjid | Tribal Area (Khyber Agency) | *Plasmodium* | *P.falciparum* | 2743 |  |
| 227 | Ali Masjid | Tribal Area (Khyber Agency) | *Plasmodium* | *P.falciparum* | 3652 |  |
| 228 | Ali Masjid | Tribal Area (Khyber Agency) | *Plasmodium* | *P.falciparum* | 3434 |  |
| 229 | Ali Masjid | Tribal Area (Khyber Agency) | *Plasmodium* | *P.falciparum* | 2443 |  |
| 230 | Ali Masjid | Tribal Area (Khyber Agency) | *Plasmodium* | *P.falciparum* | 2354 |  |
| 231 | Ali Masjid | Tribal Area (Khyber Agency) | *Plasmodium* | *P.falciparum* | 3330 |  |
| 232 | Ali Masjid | Tribal Area (Khyber Agency) | *Plasmodium* | *P.falciparum* | 1073 |  |
| 233 | Ali Masjid | Tribal Area (Khyber Agency) | *Plasmodium* | *P.falciparum* | 2679 |  |
| 234 | Ali Masjid | Tribal Area (Khyber Agency) | *Plasmodium* | *P.falciparum* | 3176 |  |
| 235 | Ali Masjid | Tribal Area (Khyber Agency) | *Plasmodium* | *P.falciparum* | 1259 |  |
| 236 | Ali Masjid | Tribal Area (Khyber Agency) | *Plasmodium* | *P.falciparum* | 1322 |  |
| 237 | Ali Masjid | Tribal Area (Khyber Agency) | *Plasmodium* | *P.falciparum* | 1863 |  |
| 238 | Ali Masjid | Tribal Area (Khyber Agency) | *Plasmodium* | *P.falciparum* | 2439 |  |
| 239 | Ali Masjid | Tribal Area (Khyber Agency) | *Plasmodium* | *P.falciparum* | 1358 |  |
| 240 | Ali Masjid | Tribal Area (Khyber Agency) | *Plasmodium* | *P.falciparum* | 2988 |  |
| 241 | Ali Masjid | Tribal Area (Khyber Agency) | *Plasmodium* | *P.falciparum* | 3402 |  |
| 242 | Ali Masjid | Tribal Area (Khyber Agency) | *Plasmodium* | *P.falciparum* | 3610 |  |
| 243 | Ali Masjid | Tribal Area (Khyber Agency) | *Plasmodium* | *P.falciparum* | 1050 |  |
| 244 | Ali Masjid | Tribal Area (Khyber Agency) | *Plasmodium* | *P.falciparum* | 1406 |  |
| 245 | Ali Masjid | Tribal Area (Khyber Agency) | *Plasmodium* | *P.falciparum* | 2563 |  |
| 246 | Ali Masjid | Tribal Area (Khyber Agency) | *Plasmodium* | *P.falciparum* | 2940 |  |
| 247 | Ali Masjid | Tribal Area (Khyber Agency) | *Plasmodium* | *P.falciparum* | 3816 |  |
| 248 | Ali Masjid | Tribal Area (Khyber Agency) | *Plasmodium* | *P.falciparum* | 2021 |  |
| 249 | Ali Masjid | Tribal Area (Khyber Agency) | *Plasmodium* | *P.falciparum* | 3545 |  |
| 250 | Ali Masjid | Tribal Area (Khyber Agency) | *Plasmodium* | *P.falciparum* | 1268 |  |
| 251 | Ali Masjid | Tribal Area (Khyber Agency) | *Plasmodium* | *P.falciparum* | 2842 |  |
| 252 | Ali Masjid | Tribal Area (Khyber Agency) | *Plasmodium* | *P.falciparum* | 1032 |  |
| 253 | Ali Masjid | Tribal Area (Khyber Agency) | *Plasmodium* | *P.falciparum* | 2507 |  |
| 254 | Ali Masjid | Tribal Area (Khyber Agency) | *Plasmodium* | *P.falciparum* | 1160 |  |
| 255 | Ali Masjid | Tribal Area (Khyber Agency) | *Plasmodium* | *P.falciparum* | 3796 |  |
| 256 | Ali Masjid | Tribal Area (Khyber Agency) | *Plasmodium* | *P.falciparum* | 2972 |  |
| 257 | Ali Masjid | Tribal Area (Khyber Agency) | *Plasmodium* | *P.falciparum* | 3231 |  |
| 258 | Ali Masjid | Tribal Area (Khyber Agency) | *Plasmodium* | *P.falciparum* | 3694 |  |
| 259 | Ali Masjid | Tribal Area (Khyber Agency) | *Plasmodium* | *P.falciparum* | 2644 |  |
| 278 | Lahore | Punjab | *Plasmodium* | *P.vivax* | 1992 |  |
| 279 | Lahore | Punjab | *Plasmodium* | *P.vivax* | 2499 |  |
| 280 | Sheikhopora | Punjab | *Plasmodium* | *P.vivax* | 1165 |  |
| 281 | Lahore | Punjab | *Plasmodium* | *P.falciparum* |  | 2202 |
| 282 | Lahore | Punjab | *Plasmodium* | *P.falciparum* |  | 3266 |
| 283 | Shanaweri Zagiri | Tribal Area (Orakzai Agency) | *Plasmodium* | *P.falciparum* |  | 2109 |
| 284 | Layyah | Punjab | *Plasmodium* | *P.vivax + P.falciparum* | 1283 | 2102 |
| 285 | Shanaweri Zagiri | Tribal Area (Orakzai Agency) | *Plasmodium* | *P.vivax + P.falciparum* | 1272 | 2588 |
| 260 | Lahore | Punjab | *Plasmodium* | Negative |  | 3155 |
| 261 | Lahore | Punjab | *Plasmodium* | Negative |  | 3201 |
| 262 | Lahore | Punjab | *Plasmodium* | Negative |  | 4209 |
| 263 | Lahore | Punjab | *Plasmodium* | Negative |  | 1358 |
| 264 | Sind | Punjab | *Plasmodium* | Negative |  | 1217 |
| 265 | DG Khan | Punjab | *Plasmodium* | Negative |  | 4328 |
| 266 | Multan | Punjab | *Plasmodium* | Negative |  | 2185 |
| 267 | Sheikhopora | Punjab | *Plasmodium* | Negative |  | 3314 |
| 268 | Lahore | Punjab | *Plasmodium* | Negative |  | 4382 |
| 269 | Lahore | Punjab | *Plasmodium* | Negative |  | 3255 |
| 270 | Lahore | Punjab | *Plasmodium* | Negative |  | 3207 |
| 271 | Sheikhupura | Punjab | *Plasmodium* | Negative |  | 2262 |
| 272 | Ali Masjid | Tribal Area (Khyber Agency) | *Plasmodium* | Negative | 1978 |  |
| 273 | Ali Masjid | Tribal Area (Khyber Agency) | *Plasmodium* | Negative | 3420 |  |
| 274 | Ali Masjid | Tribal Area (Khyber Agency) | *Plasmodium* | Negative | 1015 |  |
| 275 | Ali Masjid | Tribal Area (Khyber Agency) | *Plasmodium* | Negative | 2435 |  |
| 276 | Ali Masjid | Tribal Area (Khyber Agency) | *Plasmodium* | Negative | 2741 |  |
| 277 | Shanaweri Zagiri | Tribal Area (Orakzai Agency) | *Plasmodium* | Negative | 2120 |  |
| 286 | Sheikhupura | Punjab | *Plasmodium* | Negative | Negative | Negative |
| 287 | Kasur | Punjab | *Plasmodium* | Negative | Negative | Negative |
| 288 | Kasur | Punjab | *Plasmodium* | Negative | Negative | Negative |
| 289 | Larkana | Punjab | *Plasmodium* | Negative | Negative | Negative |
| 290 | Sheikhupura | Punjab | *Plasmodium* | Negative | Negative | Negative |
| 291 | Lahore | Punjab | *Plasmodium* | Negative | Negative | Negative |
| 292 | Sheikhupura | Punjab | *Plasmodium* | Negative | Negative | Negative |
| 293 | Lahore | Punjab | *Plasmodium* | Negative | Negative | Negative |
| 294 | Gijrat | Punjab | *Plasmodium* | Negative | Negative | Negative |
| 295 | Multan | Punjab | *Plasmodium* | Negative | Negative | Negative |
| 296 | Lahore | Punjab | *Plasmodium* | Negative | Negative | Negative |
| 297 | Sheikhupura | Punjab | *Plasmodium* | Negative | Negative | Negative |
| 298 | Sahiwal | Punjab | *Plasmodium* | Negative | Negative | Negative |
| 299 | Multan | Punjab | *Plasmodium* | Negative | Negative | Negative |
| 300 | Multan | Punjab | *Plasmodium* | Negative | Negative | Negative |
| 301 | Multan | Punjab | *Plasmodium* | Negative | Negative | Negative |
| 302 | Lahore | Punjab | *Plasmodium* | Negative | Negative | Negative |
| 303 | Lahore | Punjab | *Plasmodium* | Negative | Negative | Negative |
| 304 | Sheikhupura | Punjab | *Plasmodium* | Negative | Negative | Negative |
| 305 | Lahore | Punjab | *Plasmodium* | Negative | Negative | Negative |
| 306 | Narowal | Punjab | *Plasmodium* | Negative | Negative | Negative |
| 307 | Lahore | Punjab | *Plasmodium* | Negative | Negative | Negative |
| 308 | Okara | Punjab | *Plasmodium* | Negative | Negative | Negative |
| 309 | Larkana | Punjab | *Plasmodium* | Negative | Negative | Negative |
| 310 | Larkana | Punjab | *Plasmodium* | Negative | Negative | Negative |
| 311 | Lahore | Punjab | *Plasmodium* | Negative | Negative | Negative |
| 312 | Lahore | Punjab | *Plasmodium* | Negative | Negative | Negative |
| 313 | Lahore | Punjab | *Plasmodium* | Negative | Negative | Negative |
| 314 | Lahore | Punjab | *Plasmodium* | Negative | Negative | Negative |
| 315 | Lahore | Punjab | *Plasmodium* | Negative | Negative | Negative |
| 316 | Lahore | Punjab | *Plasmodium* | Negative | Negative | Negative |
| 317 | Lahore | Punjab | *Plasmodium* | Negative | Negative | Negative |
| 318 | Lahore | Punjab | *Plasmodium* | Negative | Negative | Negative |
| 319 | Lahore | Punjab | *Plasmodium* | Negative | Negative | Negative |
| 320 | Lahore | Punjab | *Plasmodium* | Negative | Negative | Negative |
| 321 | Lahore | Punjab | *Plasmodium* | Negative | Negative | Negative |
| 322 | Lahore | Punjab | *Plasmodium* | Negative | Negative | Negative |
| 323 | Lahore | Punjab | *Plasmodium* | Negative | Negative | Negative |
| 324 | Lahore | Punjab | *Plasmodium* | Negative | Negative | Negative |
| 325 | Lahore | Punjab | *Plasmodium* | Negative | Negative | Negative |
| 326 | Lahore | Punjab | *Plasmodium* | Negative | Negative | Negative |
| 327 | Lahore | Punjab | *Plasmodium* | Negative | Negative | Negative |
| 328 | Lahore | Punjab | *Plasmodium* | Negative | Negative | Negative |
| 329 | Lahore | Punjab | *Plasmodium* | Negative | Negative | Negative |
| 330 | Lahore | Punjab | *Plasmodium* | Negative | Negative | Negative |
| 331 | Lahore | Punjab | *Plasmodium* | Negative | Negative | Negative |
| 332 | Lahore | Punjab | *Plasmodium* | Negative | Negative | Negative |
| 333 | Lahore | Punjab | *Plasmodium* | Negative | Negative | Negative |
| 334 | Faisalabad | Punjab | *Plasmodium* | Negative | Negative | Negative |
| 335 | Lahore | Punjab | *Plasmodium* | Negative | Negative | Negative |
| 336 | Lahore | Punjab | *Plasmodium* | Negative | Negative | Negative |
| 337 | Lahore | Punjab | *Plasmodium* | Negative | Negative | Negative |
| 338 | Lahore | Punjab | *Plasmodium* | Negative | Negative | Negative |
| 339 | Okara | Punjab | *Plasmodium* | Negative | Negative | Negative |
| 340 | Lahore | Punjab | *Plasmodium* | Negative | Negative | Negative |
| 341 | Gujrat | Punjab | *Plasmodium* | Negative | Negative | Negative |
| 342 | Okara | Punjab | *Plasmodium* | Negative | Negative | Negative |
| 343 | Sheikhopora | Punjab | *Plasmodium* | Negative | Negative | Negative |
| 344 | Lahore | Punjab | *Plasmodium* | Negative | Negative | Negative |
| 345 | Lahore | Punjab | *Plasmodium* | Negative | Negative | Negative |
| 346 | Sheikhupura | Punjab | *Plasmodium* | Negative | Negative | Negative |
| 347 | Sheikhupura | Punjab | *Plasmodium* | Negative | Negative | Negative |
| 348 | Kasur | Punjab | *Plasmodium* | Negative | Negative | Negative |
| 349 | kasur | Punjab | *Plasmodium* | Negative | Negative | Negative |
| 350 | Lahore | Punjab | *Plasmodium* | Negative | Negative | Negative |
| 351 | Lahore | Punjab | *Plasmodium* | Negative | Negative | Negative |
| 352 | Lahore | Punjab | *Plasmodium* | Negative | Negative | Negative |
| 353 | Lahore | Punjab | *Plasmodium* | Negative | Negative | Negative |
| 354 | Lahore | Punjab | *Plasmodium* | Negative | Negative | Negative |
| 355 | Lahore | Punjab | *Plasmodium* | Negative | Negative | Negative |
| 356 | Sheikhupura | Punjab | *Plasmodium* | Negative | Negative | Negative |
| 357 | Lahore | Punjab | *Plasmodium* | Negative | Negative | Negative |
| 358 | Multan | Punjab | *Plasmodium* | Negative | Negative | Negative |
| 359 | Lahore | Punjab | *Plasmodium* | Negative | Negative | Negative |
| 360 | Shanaweri Zagiri | Tribal Area (Orakzai Agency) | *Plasmodium* | Negative | Negative | Negative |
| 361 | Shanaweri Zagiri | Tribal Area (Orakzai Agency) | *Plasmodium* | Negative | Negative | Negative |
| 362 | Shanaweri Zagiri | Tribal Area (Orakzai Agency) | *Plasmodium* | Negative | Negative | Negative |
| 363 | Shanaweri Zagiri | Tribal Area (Orakzai Agency) | *Plasmodium* | Negative | Negative | Negative |
| 364 | Ali Masjid | Tribal Area (Khyber Agency) | *Plasmodium* | Negative | Negative | Negative |
| 365 | Ali Masjid | Tribal Area (Khyber Agency) | *Plasmodium* | Negative | Negative | Negative |
